# A potential link between tuberculosis and lung cancer through non-coding RNAs

**DOI:** 10.1101/188375

**Authors:** Debmalya Barh, Sandeep Tiwari, Ranjith N. Kumavath, Vasco Azevedo

## Abstract

Pulmonary tuberculosis caused by *Mycobacterium* and lung cancer are two major causes of deaths worldwide and the former increases the risk of developing lung cancer. However, the precise molecular mechanism of *Mycobacterium* associated increased risk of lung cancer is not entirely understood. Here, using *in silico* approaches, we show that hsa-mir-21 and *M. tuberculosis* sRNA_1096 and sRNA_1414 could play important roles in the pathogenesis of both these diseases. Further, we postulated a “*Genetic remittance*” hypothesis where these sRNAs may play important roles. The sRNA_1096 could be involved in tuberculosis through multiple infectious processes, and if transferred to the host, it may activate the TLR8 mediated pro-metastatic inflammatory pathway by acting as a ligand to TLR8 similar to the mir-21 leading to lung tumorigenesis and chemo-resistance. Analogous to SH3GL1, it may also regulate cell cycle. On the other hand, sRNA_1414 is probably involved in survivability and drug response of the pathogen. However, it may be a metastatic factor for lung cancer providing EPS8L1 and SORBS1 like functions upon remittance. Further, all these three non-coding RNAs are predicted to act in rifampicin resistance in *Mycobacterium.* Currently, we are applying robust bioinformatics strategies and conducting experimental validations to confirm our *in-silico* findings and hypothesis.

## INTRODUCTION

Viral involvements and their causal roles in oncology are well accepted for various cancers including ovarian neoplasms [1], hepatocellular carcinoma [2], and lung cancer [3] among others. Although, bacterial infections are not considered as major threats to cancer, yet a number of bacterial pathogens are reported to be associated with several cancers. Some examples include: *Mycoplasma* in prostate malignancy [4], *Robinsoniella* in pancreatic cancer [5], *S. typhi, H. bilis, H. hepaticus*, and *E. coli* in carcinoma of the gallbladder [6], *Chlamydia* in cervical cancer [7], and *Mycobacterium* in lung cancer[8-14].

Lung cancer is the leading cause of all cancer related deaths with recently estimated 1.6 million deaths worldwide [15]. Similar to lung cancer, pulmonary tuberculosis caused by *M. tuberculosis* is a global health problem. It is one of the major causes of death amongst infectious diseases and according to WHO 2013 report, it is estimated that 9 million people are infected and 1.5 million are died from tuberculosis in 2012 [16]. Several reports have documented co-existence of tuberculosis and lung cancer [14,17-20] and pulmonary tuberculosis is a risk factor for developing lung cancer [17-20]. However, it is not yet fully established at molecular level, how the *Mycobacterium* increases susceptibility to lung cancer. Some report say that *M. tuberculosis* induces ROS mediated DNA damage pathway and produces epiregulin growth factor to induce cell proliferation [21]; while other study indicates mechanisms along with COX-2 medicated activation of inflammatory pathway in *M. tuberculosis* associated carcinogenesis [22].

Bacterial small regulatory RNAs (sRNAs) are a class of small non-coding RNAs of 40-500 nt in length that regulate various essential patho-physiologies in bacteria such as outer membrane protein biogenesis, virulence, quorum sensing etc. sRNA functions through complementary base-pairing with 3′‐ or 5′-UTRs of target mRNAs to inhibit translation, alters activity of a protein by directly binding, and by mimicking RNA and DNA structures [23-25]. Several sRNAs have been identified or predicted from *M. tuberculosis* [26-28] having probable role in pulmonary tuberculosis pathogenesis [26,29,30]. However, no report so far is available on sRNA from *M. tuberculosis* having role in lung cancer. On the other hand, human micro RNAs (miRNAs) are small non-coding RNAs of 20-25 nt length that inhibit post-transcriptional gene regulation by complementary base pairing at the 3′ ‐UTRs of target mRNAs and regulate various patho-physiological conditions including cell cycle regulation, cell differentiation, development, metabolism, aging, different types of cancers, metabolic disorders, and neuronal diseases etc. [31,32]. Several miRNAs have been implemented to be associated with lung cancer having causative roles and diagnostic potentials. Similarly, a number of miRNAs are found deregulated in pulmonary tuberculosis patients [33-35].

Since the sRNAs and miRNAs are similar in structure, their mode of actions, and the miRNAs are associated with both pulmonary tuberculosis and lung cancer; we hypothesized that *Mycobacterium* sRNAs may be associated with lung cancer tumorigenesis. Since *Mycobacterium* infection is a risk factor in developing lung cancer, here we postulated a “*Genetic remittance*” hypothesis which presumes that *M. tuberculosis* sRNAs having similarity with human miRNAs (that are associated with either pulmonary tuberculosis or lung cancer or both these two diseases) are transmitted to the host during *Mycobacterium* infection, remained within the human, and act as a predisposition factor to increase the risk of lung cancer.

In this study, using *in silico* strategies, we aimed to understand (i) the role of *M. tuberculosis* sRNAs in lung carcinogenesis; (ii) if there are structural and functional similar *M. tuberculosis* sRNAs to human miRNAs responsible for pathogenesis in both pulmonary tuberculosis and lung cancer and their probable functions in these diseases; and (iii) proof of concept of our “*Genetic remittance*” hypothesis.

## METHODS

### Collection of human miRNAs and M. tuberculosis siRNAs

We used PubMed (http://www.ncbi.nlm.nih.gov/pubmed), miRegulome [32], miR2Disease [36], TUMIR [37], sRNAdb [38], and BSRD [39] databases to get the data. The data deposited in these databases during January 2006 to November 2015 were searched. In the first step, we collected all the validated deregulated miRNAs associated with lung cancer and pulmonary tuberculosis from published literature indexed in PubMed (http://www.ncbi.nlm.nih.gov/pubmed) and from databases such as miRegulome [32], miR2Disease [36], and TUMIR [37]. The common miRNAs associated with both the diseases were manually identified and listed out separately. Similar to miRNA, we collected all reported and novel *M. tuberculosis* sRNAs by means of PubMed literature mining and searching sRNA databases such as sRNAdb [38] and BSRD [39]. For PubMed search, specific key words such as “miRNA + *M. tuberculosis*”, “miRNA + tuberculosis”, “miRNA + lung cncer”, sRNA + *M. tuberculosis”, “*sRNA + tuberculosis” etc. were used.

### Prediction of miRNAs that may function as sRNA or vice versa

To achieve this, we used a simple strategy. Since the sRNAs and miRNAs are of very short in sequence, we presumed that small sequence motifs in these non-coding RNAs are very important for their specific functions. Therefore, we used comparative BLASTn (miRNA against sRNA) with default parameters in order to identify if there is any human miRNA (from the group of miRNAs common to both lung cancer and pulmonary tuberculosis) having sequence similarity to any *M. tuberculosis* sRNA, so that, they may function similarly. From the BLASTn results, the specific motif sequences that are common in sRNAs and miRNAs were noted down.

### Functional annotation of M. tuberculosis sRNAs

The functional annotation of the sRNAs was carried out using target based reverse annotation approaches following a modified protocol as described by Barh et al., 2013 [40,41]. In brief, we identified the validated targets of sRNA using sRNATarBase [42] database and RNApredator [43] was used to predict putative targets in *M. tuberculosis*, H37Rv. Top 100 targets were used for functional annotation of using DAVID functional annotation tool [44]. Further, we presumed that, if there is a coding gene that has identity with sRNA; the function of the sRNA could be similar to that gene. Therefore, we performed sRNA BLASTn against *M. tuberculosis* genome using NCBI BLASTn server (http://blast.ncbi.nlm.nih.gov/Blast.cgi) to identify if there is any coding sequence present in *M. tuberculosis* genome having sequence identity with the sRNA. To check if the sRNA is targeting an essential gene of *M. tuberculosis*, H37Rv, we used Database of Essential Genes (DEG) [45] BLASTn and to check if the target could be a drug target, we used the strategy as described by Barh et al., 2013 [41]. Further, we did miRNA (that is having sequence similarity with sRNA) BLASTn *M. tuberculosis* genome in NCBI BLASTn server with default parameters to check if any *M. tuberculosis* coding sequence is matching with the miRNA sequence. Since, in this against way identified miRNA shares sRNA sequence, if there is a coding sequence that matches with the miRNA; we may assume that, the matched coding sequence of the sRNA and the miRNA may have similar function.

### Testing “Genetic remittance” hypothesis

To test our *”Genetic remittance”* hypothesis, we performed BLASTn of the *M. tuberculosis* sRNAs having sequence similarity with human miRNA (that are associated with both pulmonary tuberculosis and lung cancer) against the human genome on NCBI BLASTn server in order to identify if there is any human coding sequence having identity with the sRNAs. If there is a sequence match and that human gene sequence is responsible for any tumorigenesis process, we assumed that, there may be possibility that these sRNAs may transfer to the host during *Mycobacterial* infection, and with a yet unknown mechanism regulate those human genes to increase the risk of lung tumorigenesis. Further, as those sRNAs are sharing human miRNA that are associated with lung cancer, there could be possibility that the predisposition of sRNA may have synergetic effects in presence of miRNA that leads to lung cancer.

## RESULTS

### Common miRNAs in pulmonary tuberculosis and lung cancer

From various literature and databases, we collected differential expression of 186 human miRNAs in pulmonary tuberculosis and 242 miRNAs in lung cancer patients. However, while we checked for the common miRNAs that are deregulated both in pulmonary tuberculosis and lung cancer, we found the number is only 45 (**Supplementary Table S1**).

### M. tuberculosis sRNA_1096 and sRNA_1414 shares hsa-mir-21 sequence

Form the sRNAdb [38] and BSRD [39] databases and literature [27,46] we collected 120 reported *M. tuberculosis* sRNAs and their sequences. The comparative BLASTn between the 120 sRNAs and the common 45 human miRNAs in pulmonary tuberculosis and lung cancer revealed that the mature human miRNA hsa-mir-21-5p having very short sequence similarity with *M. tuberculosis* sRNA_1096 and sRNA_1414. Both these sRNAs are experimentally validated in *M. tuberculosis* [27]. Human hsa-mir-21-5p and *M. tuberculosis* sRNA_1096 share a common sequence of GTTG/ GUUG, while the common sequence between hsa-mir-21-5p and *M. tuberculosis* sRNA_1414 is ATCAG/ AUCAG (**Supplementary Table S2**). ***Functional annotation of sRNA_1096 and sRNA_1414***

To understand the functions of sRNA_1096 and sRNA_1414 in *M. tuberculosis*, first we used sRNATarBase [42] to search validated targets of these sRNAs. However, we did not get any target from this database. Therefore, we used RNApredator [43]to predict the targets of these sRNAs. The top 100 targets based on the Z-score of RNApredator were further used for DAVID functional annotation.

For sRNA_1096, phosphate-binding protein pstS 2 (Z-score: ‐11.29) was found to be the best target. pstS2 is an component system and ABC transporters pathways. Further, among the top 100 targets, targets are PE PGRS family proteins (**Supplementary Table S3**). DAVID functional annotation analysis of the top 100 targets of sRNA_1096 shows that most targets are membrane located and the two-component system is the top annotation cluster.

The top target of sRNA_1414 is found to be glycyl-tRNA synthetase / glycine‐‐tRNA ligase (glyS) (Z-score: ‐11.41) by RNApredator [43] (**Supplementary Table S3**). The drug transporter activity is ranked as the first annotation cluster and most targets are transmembrane proteins as observed through DAVID functional annotation analysis. According to the Database of Essential Genes (DEG) [45], the *M. tuberculosis* glyS is an inorganic phosphate transmembrane transporter that is involved in two-fourteen essential gene in the pathogen but has 31% identity at protein level with human glycine‐‐ tRNA ligase as per NCBI human BLASTp.

When we performed sRNA_1096 BLASTn against *M. tuberculosis* genome, a very short similarity “CCGTCACCGTTG” was observed with *M. tuberculosis* Arabinosyltransferase *EmbC* (*embC*) gene The BLASTn of sRNA_1414 with *M. tuberculosis* genome did not show any hit with any specific protein coding gene of the pathogen.

### Common function of sRNA_1096, sRNA_1414, and hsa-mir-21

As in previous analysis we found there are sequence similarities among hsa-mir-21 and *M. tuberculosis* sRNA_1096 and sRNA_1414; we performed a NCBI BLASTn of hsa-mir-21-5p against the *M. tuberculosis* genome. We observed that the *M. tuberculosis* rpoB gene that provides rifampin/ rifampicin resistant in *M. tuberculosis* has sequence similarity with mir-21 and having both the GTTG and ATCAG short-stretch sequences that are present in *M. tuberculosis* sRNA_1096 and sRNA_1414, respectively. Therefore, all these non-coding may be involved in rifampicin resistance in pulmonary tuberculosis.

### Support to “Genetic remittance” hypothesis

To support our “*Genetic remittance*” hypothesis, we tried to identify if there is any sequence match of sRNA_1096 and sRNA_1414 in human coding sequence. The BLASTn of sRNA_1096 against human genome hits with SH3GL1 (SH3-domain GRB2-like 1) and some sequence of sRNA_1414 matches with human EPS8L1 (EPS8-like 1) and SORBS1 (Sorbin and SH3 domain containing 1). All these human genes are associated with tumorigenesis, thus providing a preliminary support to our hypothesis.

## DISCUSSION

In this study, we found that there could be correlations between lung cancer and pulmonary tuberculosis at non-coding RNA level. *M. tuberculosis* sRNA_1096 and sRNA_1414, and human hsa-mir-21-5p could probably are the links to explain why the pulmonary tuberculosis is a risk factor in developing lung cancer. Among the 45 human miRNAs that are deregulated both in both the diseases (**Supplementary Table S1**) and 120 reported *M. tuberculosis* sRNAs is genome, we found that there are sequence similarities among human hsa-mir-21-5p and *M. tuberculosis* sRNA_1096 and sRNA_1414 (**Supplementary Table S2**).

The onco-miR hsa-mir-21 is frequently upregulated in lung cancer [47-55] while it is downregulated in CD4^+^ T cells in tuberculosis patients [56]. However, hsa-mir-21 is found upregulated in the host during *M. bovis* BCG infection [57] (**Supplementary Table S1**). It is also observed that mir-21 plays a role in T-cell immunity against *M. tuberculosis*is [56] and found to affect the anti-mycobacterial T‐ cell response through targeting IL12 and BCL2 [57].

Reports suggest that the bacteria induced carcinogenesis occurred in multiple ways. These mechanisms include induction or interference of chronic inflammatory and other signalling cascades including TLRs (Toll-like receptors) signalling and acetaldehyde metabolism pathways in various cancers [58-60]. TLRs signalling are generally involved in innate and adaptive immune responses. However, activation of TLRs signalling promotes tumor cell proliferation, growth, invasion, and metastasis [61]. In lung cancer, activation of TLR7 and TLR8 increases survival and chemo-resistance of the tumor cells, therefore these two TLRs could be targets for tumor immunotherapy [62]. In *Mycobacterial* infection, TLRs signalling regulates host innate and inflammatory responses and determines the disease outcome [63]. Polymorphisms and over expression of TLR8 is associated with pulmonary tuberculosis susceptibility and infection, respectively [64]. However, the precise mechanism of TLR8 in pulmonary tuberculosis is not yet known [65].

Fabbri *et al* in 2012 first reported the novel mechanism of onco-miR mir-21 that can act as a ligand for TLR8 to induce inflammatory response leading to tumor growth and metastasis [66]. This study also showed that the “GUUG” motif miR-21 directly binds to TLR8 to induce the TLR-mediated pro-metastatic inflammatory response.

### M. tuberculosis sRNA_1096

In our analysis, we found that the GTTG/ GUUG sequence is conserved in hsa-miR-21 and experimentally validated *M. tuberculosis* sRNA_1096 (**Supplementary Table S2**). Therefore, it may be implemented that the *M. tuberculosis* sRNA_1096 may bind to TLR8 in a similar way the miR-21 does through its GUUG motif and thus may activate immune and inflammatory responses and explains a role of TLR8 in pulmonary tuberculosis.

Further, as per our analysis, *M. tuberculosis* sRNA_1096 may play a role in two-component system pathway and may regulate PE PGRS family and membrane located proteins. PE PGRS family proteins act as variable surface antigens and are involved in multiple levels of the infectious process, modulation of innate immune responses [67], and virulence [68] in *M. tuberculosis*. Similarly, the two-component system is crucial in *Mycobacterial* survival and pathogenicity [69]. In *M. tuberculosis* genome, “CCGTCACCGTTG” sequence of the sRNA_1096 is present in Arabinosyltransferase EmbC (embC) gene which is involved in bacteria-host interactions and also modulates immune response in *M. tuberculosis* [70]. Therefore, *M. tuberculosis* sRNA_1096 could play an important role in *M. tuberculosis* pathogenesis leading to pulmonary tuberculosis.

In support to our “*Genetic remittance*” hypothesis, we predicted that if the *M. tuberculosis* sRNA_1096 is present or predisposes to human, it may modulate human SH3GL1 (SH3-domain GRB2-like 1) which plays a role in endocytosis [71], regulation of cell cycle in leukaemia [72], positive regulation of cell proliferation and inhibition of apoptosis in multiple myeloma [73], and oncogenesis in gliomas [74]. Hence, sRNA_1096 may play a critical role in lung cancer risk. Further, if the sRNA_1096 acts as a ligand to TLR8 similar to mir-21, upon remittance to the host, it may activate TLR8 pathway and along with increasing risk it may also regulate chemo-resistance [62] in lung cancer individually or in combination with mir-21.

Therefore, in summary, we postulate that, the *M. tuberculosis* sRNA_1096 is involved in pulmonary tuberculosis pathogenesis through multiple infectious processes including TLR8 mediated pathway. The sRNA_1096 may be transported to host and predisposed during *M. tuberculosis* infection and later acts as ligand to TLR8 through its GTTG/ GUUG sequence similar to mir-21 to activate TLR8 mediated pro-metastatic pathways and chemo-resistance in lung cancer. Similar to SH3GL1, it may also regulate tumorigenic inflammatory response and cell cycle, respectively leading to lung cancer. Hence, sRNA_1096 may be an emerging marker for tuberculosis and lung cancer risk and chemo-resistance in combination with mir-21.

### M. tuberculosis sRNA_1414

On the other hand, we observed ATCAG/ AUCAG as the common sequence between hsamir-21-5p and *M. tuberculosis* sRNA_1414 (**Supplementary Table S2**). The sRNA_1414 is predicted to target glycyl-tRNA synthetase (glyS) which is an essential gene in *M. tuberculosis*. Further, we observed that sRNA_1414 may regulate drug transporter activity. Aspartyl-tRNA synthetase and tyrosyl-tRNA synthetase are important drug targets in *M. tuberculosis,* [75,76] and polymorphisms in aspartyl-tRNA synthetase is associated with drug resistance mechanism in this pathogen [77]. Although, we predicted glyS is as an essential gene in *M. tuberculosis,* being a human homolog, it is not a suitable target. Thus sRNA_1414 probably be involved in regulating the survivability and drug response of the pathogen.

To evaluate our “*Genetic remittance*” hypothesis, while we did the BLASTn of sRNA_1414 against human genome, we found sequence matches with human EPS8L1 (EPS8-like 1) and SORBS1 (Sorbin and SH3 domain containing 1). EPS8L1 encodes a protein that is related to epidermal growth factor receptor pathway substrate 8 and is involved in regulation of Rho protein signal transduction [78] which is associated with small cell lung cancer migration [79]. Similarly, SORBS1 (Sorbin and SH3 domain containing 1) plays an important role in cell-matrix adhesion [80] a key process in cell migration. Therefore, if the *M. tuberculosis* sRNA_1414 is transferred to the host during the infection, it may lead to lung cancer metastasis in later stage functioning similar to EPS8L1 and SORBS1.

### Common functions of M. tuberculosis sRNA_1096 and sRNA_1414

Since the hsa-mir-21 sequence GTTG /GUUA is shared by sRNA_1096 and ATCAG/ AUCAG by sRNA_1414 (**Supplementary Table S2**), we tried to predict what could common role of these non-coding RNAs. Our hsa-mir-21 BLASTn against *M. tuberculosis* shows that these GTTG /GUUA and ATCAG/ AUCAG sequences are present in *M. tuberculosis rpoB* gene that provides rifampin/resistant in *M. tuberculosis* [81-84]. Therefore, we presume that all these non-coding RNAs: hsa-mir-21, sRNA_1096, and sRNA_1414 could be involved in rifampicin resistance and an up regulation of mir-21 in tuberculosis patient may be a marker of rifampicin resistance.

## CONCLUSIONS

In this work, we hypothesise that *M. tuberculosis* sRNA_1096 and sRNA_1414 may be involved in increased risk of developing lung cancer metastasis and chemo-resistance provided they are transferred to human and during *Mycobacterial* infection. These sRNAs may have synergetic effects in presence of human mir-21 that is associated with lung cancer. This analysis also provides an insight on why the *Mycobacterial* infection increases the risk of developing lung cancer and the possible TLR8 mediated molecular mechanisms. Further, all these non-coding RNAs could be involved in rifampicin resistance in *Mycobacterium*. Therefore, all these non-coding RNAs are potential biomarkers for assessing rifampicin resistance and lung cancer risk in pulmonary tuberculosis patients. A robust bioinformatics approach as outlined in Figure-1 including domain/motif analysis along with experimental validations are required to establish this proof of concept.

**Figure 1.**
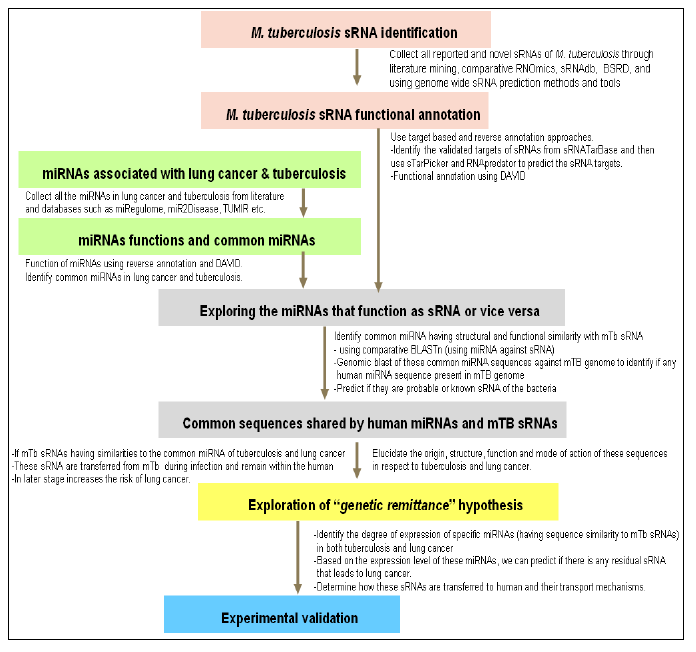
The integrated bioinformatics and experimental validation strategy to elucidate the relationship between tuberculosis and lung cancer linked through non-coding RNAs.

## Acknowledgments

Sandeep Tiwari acknowledges the “TWAS-CNPq Postgraduate Fellowship Programme” for granting a fellowship for doctoral studies. Debmalya Barh acknowledges the “TWAS-CNPq Postdoctoral Fellowship Programme” for granting a fellowship for postdoctoral studies.

## Author Contributions

Conceived, designed the experiment, collected and analysed initial data, coordinated and led the entire project: DB, performed all *in-silico* analysis: DB, ST. Cross-checked all data, analysis: RNK, VA. Wrote the paper: DB. All authors read and approved the manuscript

## Supplementary table legends

**Supplementary Table S1**

Common miRNAs that are deregulated both in pulmonary tuberculosis and lung cancer.

**Supplementary Table S2**

Sequences and similarities among of *M. tuberculosis* sRNA_1096, sRNA_1414, and human hsa-miR-21.

**Supplementary Table S3**

RNApredator based predicted targets of *M. tuberculosis* sRNA_1096 and sRNA_1414.

